# Regular geometry and hexagonal structure of honeycomb results from the optimisation of cylindrical cells built in close proximity

**DOI:** 10.1101/2022.07.13.499872

**Authors:** Vincent Gallo, Alice D. Bridges, Joseph L. Woodgate, Lars Chittka

## Abstract

The hexagonal structure of honeycomb maximises storage volume while minimising the amount of wax required for its construction. How honeybee builders achieve this geometry, however, remains unclear. Previously, our group identified behavioural patterns that were triggered in builders when they encountered certain sub-scale features associated with partially constructed comb, which resulted in the alignment of new cells to small concavities and the construction of cell walls between two of these stimuli. This caused new cells to be built in the proper locations without the need for explicit instructions . Here, we investigated whether the hexagonal geometry of honeycomb cells resulted from the dense packing of cells that would otherwise have been circular tubes. We hypothesised that the reaction of a builder to a cell that is not fully enclosed by other cells would be an attempt to maximise the internal space by excavating and re-forming the surrounding walls to create a cylindrical interior.

However, the creation of a cylindrical cell would be thwarted by the activities of workers within adjacent cells also acting according to these rules. Eventually an equilibrium will emerge with walls that meet at a junction arranged so that the available angular range (360°) is sub- divided equally between the cells that meet at the junction (typically, internal angles of 120° when three cells meet). To test this hypothesis, we offered wax stimuli to comb-building honeybees, shaped to encourage or to constrain the construction of comb cells, recording the bees’ progress. We found that at an early stage cells could be an irregular shape with curved walls and unequal wall lengths and corner angles, however, when allowed further time and unconstrained access the workers reshaped the cells achieving significantly greater regularity.

## Introduction

Honeybee comb has long been the subject of human fascination, with the first recorded analysis of its structure dating back to 320 CE (Heath 1921). Honeycomb is characterised by both its regularity and its structural efficiency: a double-sided sheet of tessellated hexagonal cells that both minimises the wax required for construction and maximises storage capacity (Graham 1993; Hepburn et al. 2014; Gallo and Chittka 2018). The cells have pyramidal bases that interlock along a shared backplane, resulting in an offset of half a cell between the two sides. Thus, while the hexagonal face of a sheet of comb is characterized by three-way cell intersections, with 120° wall junctions equally spaced in two dimensions around 360°, the interface between the two sides of comb involves *four*-way cell intersections involving a cell from one face centred at the intersection of three cells from the other. At each four-way intersection, the dihedral angles are also 120°, equally spaced around 360° in a three-dimensional layout. Proper tessellation of cells along a single side of comb appears to require the placement of new cells between two extant cells, but as comb is double-sided, this actually requires new cells to be placed exactly between *three* extant cells. Honeybees seem to be particularly adept at building comb with even distributions of cell walls around intersections but how they achieve this remains to be fully understood.

Previously, our group investigated the earliest stage of comb construction: the initial placement of new cells, and how bees decide this. When offered wax stimuli that included a shallow depression, mimicking the naturally occurring clefts between two partially formed cells, workers were induced to enlarge the depression and deposit wax at its edges (Gallo et al. 2022). The workers formed cells and walls at locations which ensured proper comb tessellation in response to the existing form of the workpiece. Guided in this way, a mechanism described as stigmergy, multiple individuals are able to contribute to the overall structure of comb by reacting only to the perceived conditions in their own immediate locale.

Stigmergy, a form of self-organisation where individual participants react in response to local conditions, has previously been hypothesized to underly the coordination of activities in social insects (Bonabeau et al. 1999; Collignon and Detrain 2019). Both the parallel blades of comb formed by honeybees and the pattern of cell use within a hive were previously thought to be driven by stigmergy (Hepburn and Whiffler 1991; Theraulaz and Bonabeau 1999), and our findings provided the first evidence to suggest that stigmergy might underlie the construction of individual cells within honeycomb. In the present study, we sought to build upon these results and investigate whether stigmergy might be involved in other elements, at later stages, of comb construction.

It has been noted previously that at an early stage of honeycomb construction the cell walls are curved and the cells are approximately circular, but as construction of the comb continues, the walls become straighter and the cells more hexagonal during construction (Pirk et al. 2004; Karihaloo et al. 2013; Hepburn et al. 2014, p. 248) . While these authors described the change from curved to straight walled cells, they claimed the mechanism to be thermoplasticity of the wax but this has been rebuffed due to the wax not achieving the required temperature (Humphrey and Dykes 2008; Bauer and Bienefeld 2012). Here, we investigated whether this progression of walls from curved to straight, and of cells from circular to hexagonal cess, as well as the equalisation of angles around cell junctions, was governed by some rule-based behaviour by the bees. We posit that honeybee construction workers, if building an isolated cell, would not construct a hexagonal prism with straight walls and a pyramidal base, rather they would create one as a circular tube with hemispherical base similar to the curved, ovoid cells formed within the ground by species of solitary bees and within nest cavities by bumble bees (Goulson 2000; Danforth et al. 2019). However, honeybee cells are not isolated but are densely packed with each cell in contact with adjacent ones and the ultimate shape therefore results from equilibrium between the efforts of builders attending to each cell. That honeycomb geometry results from the compaction of cells that otherwise would have been curved, has been proposed previously (Kepler 1611; Buffon 1740; Darwin 1859; Armbruster 1920; Darchen 1959), conjectures from which we have derived our hypothesis leading to predictions that we have now tested.

Our hypothesis is that construction workers will endeavour to form a cylindrical or ovoid cell, establishing the perimeter of the cell at an ideal radius. Where such an attempt interacts with others building adjacent cells the initial conjunction will be uncoordinated and thus asymmetric but, after further work, the eventual result will be a balanced distribution of space between the conjoined cells. Cell walls within completed comb typically have a cell to either side and therefore will have been subject to the same influence exerted from both cells and equilibrium in this case would be a flat wall. Junctions where walls meet can arise through contact between three cells, and occasionally four or more (Hepburn and Whiffler 1991; Smith et al. 2021). Where the construction of a cell influences, and is influenced by, the construction of an adjacent cell, the walls meeting at the resulting junction will be arranged such that the available angular range is divided equally between all cells that share the junction.

We empirically tested predictions based on our hypothesis by offering honeybees atypical workpiece conditions in the form of specially designed wax stimuli, which they were allowed to build upon. The resulting construction was observed at an early stage, and again once further work had been undertaken. Our predictions were:-

Prediction 1 – that the internal angle between side-walls of a cell is not necessarily 120, rather that at a junction will be the available divided by the number of cells at the junction. Three walls meeting at a point will yield 360° divided by three, or 120° whereas a constrained space of 180° where two cells meet will settle as 90° for each cell.

Prediction 2 – that the angle between the base of a cell and its side walls is not necessarily 120°, rather that where cells lie opposite each other, and the mutual base will form a single face at 90° to the side wall for each cell.

Prediction 3 – cells, initially built unevenly, will be optimised by further worker activity. Regularity metrics including the area occupied by each cell, the distribution of corner angles and length of cell-walls will be improved .

## Materials and methods

### Hives

These experiments were conducted during May, June and July 2020 also June and July 2021 in England (Reigate, 51.23°N, 0.19°W). Three colonies of honeybees (*Apis mellifera*) were used, each headed by a locally reared queen housed in Modified British National hives with open mesh floors and a single brood box containing 10 conventional frames plus one to carry the experimental stimuli The frames were set transverse to the entrance, ‘warm’ alignment and the test frame positioned at, or close to, the edge of the brood area, as the 7th frame from the front. The hives were provisioned with ad libitum 1:1 sucrose solution (1.0 kg cane sugar in 1.0 l water.

### Preparation of stimuli

We created three forms of wax stimuli to investigate angular and area equalisation under different conditions (Fig. 1 a). The stimuli were fabricated using wax that had been recovered from hives within the same Reigate apiary. Flat sheets of wax, used to form the stimuli, were formed by dipping into molten wax to coat one face of a flat wooden form (75 x 40 mm). The thickness of the wax sheets could be varied by altering the number of immersions: three immersions yielded sheets of 0.5 mm-0.6 mm thickness, while six produced sheets of 1.0 mm-1.2 mm. Each sheets was cut into three pieces (25 x 40 mm), henceforth referred to as tabs, which were bonded vertically using molten wax the top bar of an otherwise empty test frame ready for placement in a hive.

**Fig. 1.**
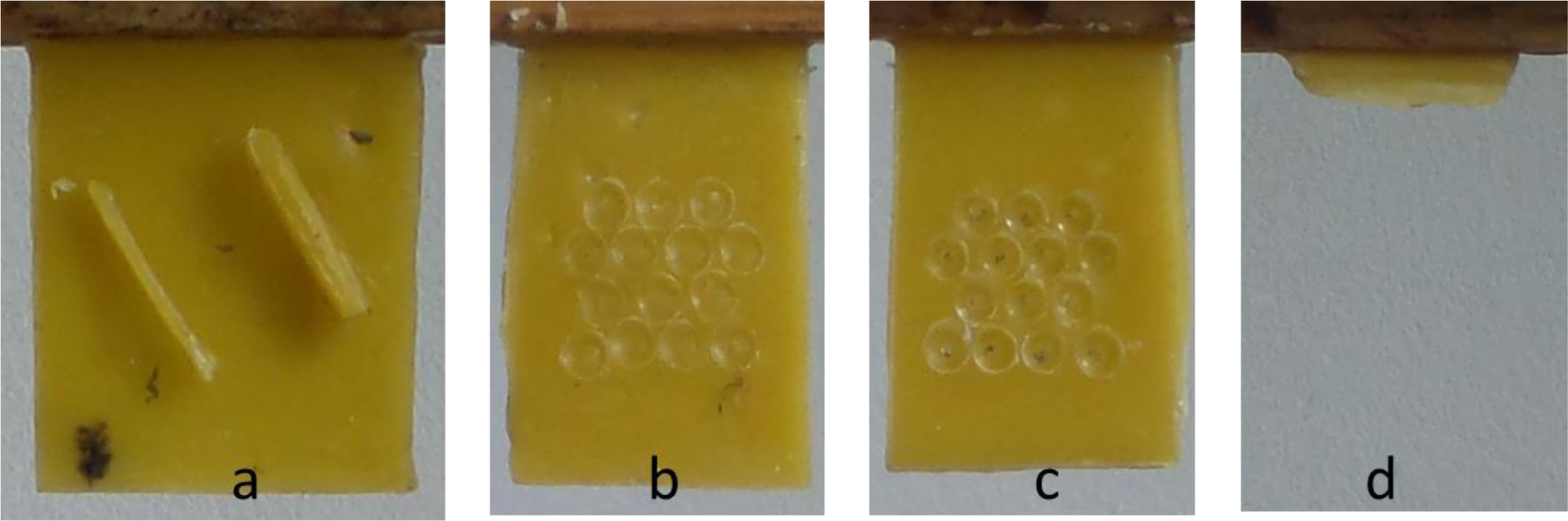
Tabs illustrating examples of the stimuli used in each experiment. These shapes were placed within the hives to allow comb to be built upon them. (a) Linear barrier of wax welded to the wax tab, used for experiment 1. (b & c) Pits, or depressions, pressed into the wax tab arranged to align pits on the front (b) and back (c), used for experiment 2. (d) Example seed strip used to encourage the construction of a independent tongues of comb, used for experiment 3.

#### Experiment 1

Our first prediction states that, the distribution of available angular space will be distributed equally between adjacent cells and redistributed between such cells as conditions change. This prediction concerning the internal angles between side-walls of cells, was tested using stimuli that each comprised a barrier, 2 mm to 3 mm in height by ∼0.6 mm thick, mounted by welding onto a wax sheet approximately 25 mm by 40 mm and 0.6 mm thick (Fig. 1 a and Fig. 2)). Our prediction was that the bees would build cells against the barrier and the barrier would constrain the available angular range to 180° thus an inter-cell wall would be aligned at 90° to the barrier (Fig. 2 c). After further construction, cells would be built on the other side of the barrier and the barrier would be eroded until the work on one side of the barrier would influence that on the other. At that, subsequent, stage the angular range around the junction would be 360° to be divided equally between three co-adjacent cells thus each cell would be adjusted to have an internal angle of 120° (Fig. 2 e).

**Fig. 2.**
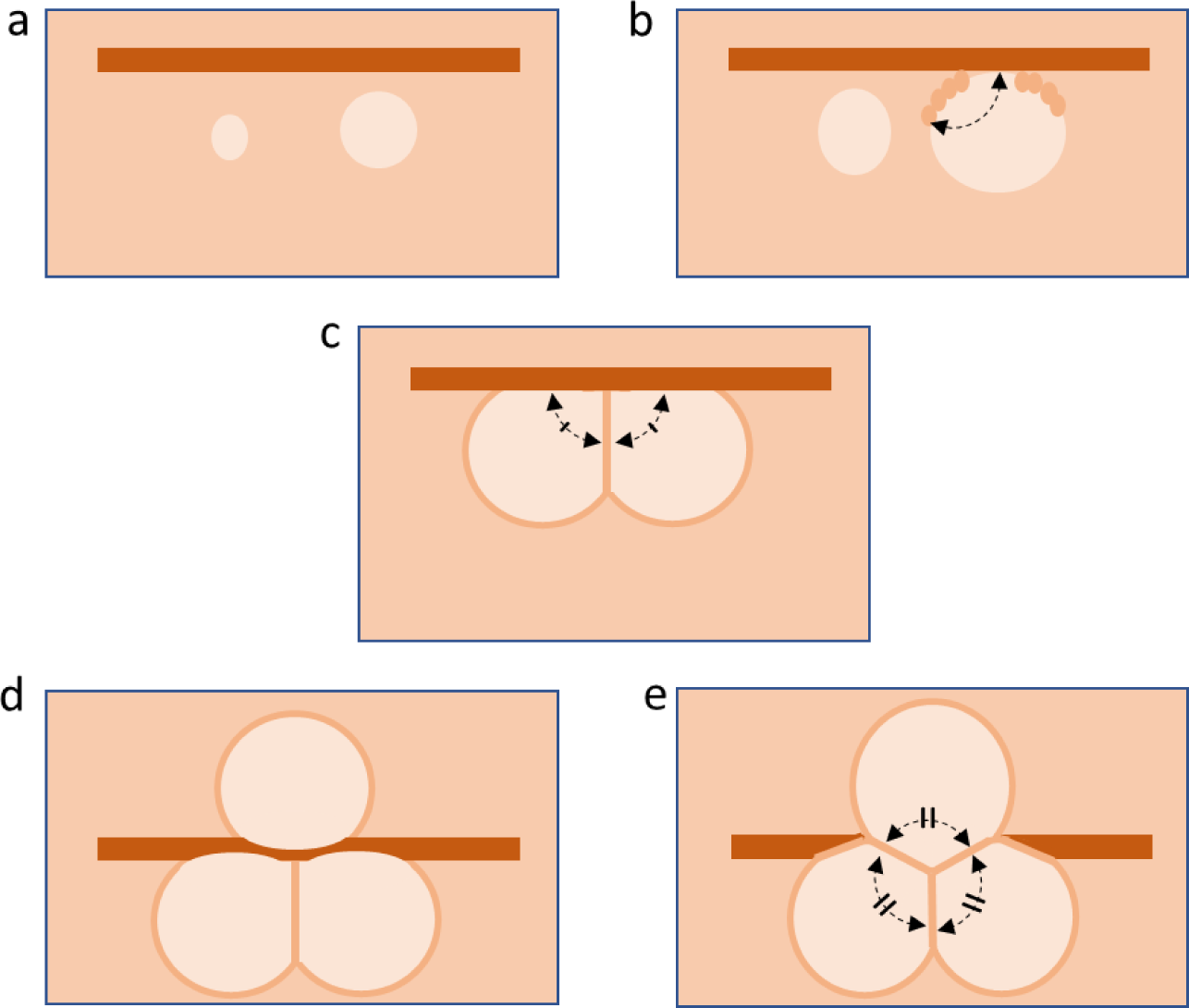
Predicted progression of cells built against a barrier. (a) Initial construction close to the barrier that will grow until a cell is built, exploiting the barrier as one of its walls. (b) the early-stage cell with peripheral deposits that form the basis of subsequent walls (c) As construction continues, a second cell built adjacent to the first will influence the mutual wall, dividing the available angle of 180° into two equal internal angles of 90°.(d) After further construction on the opposite side of the barrier, the barrier will be eroded. (e) the 3-way influence will divide the range, a full circle, between three internal angles.

#### Experiment 2

Here we tested our second prediction that the distribution of available angular space will be distributed equally between adjacent cells where the base of the cells forms the common wall. Stimuli to test this prediction comprised shallow indentations that were pressed into each side of a tab using a 4 mm-diameter domed rod forming indentations were ∼0.25 mm deep and between 3-4 mm in diameter. The depressions were pressed into both faces of the wax, arranged as an approximation of the layout and spacing of worker cells and aligned so that those on one face were each opposite one on the other face to encourage the construction of cells in direct opposition (Fig. 1 b & c and Fig. 3).

**Fig. 3.**
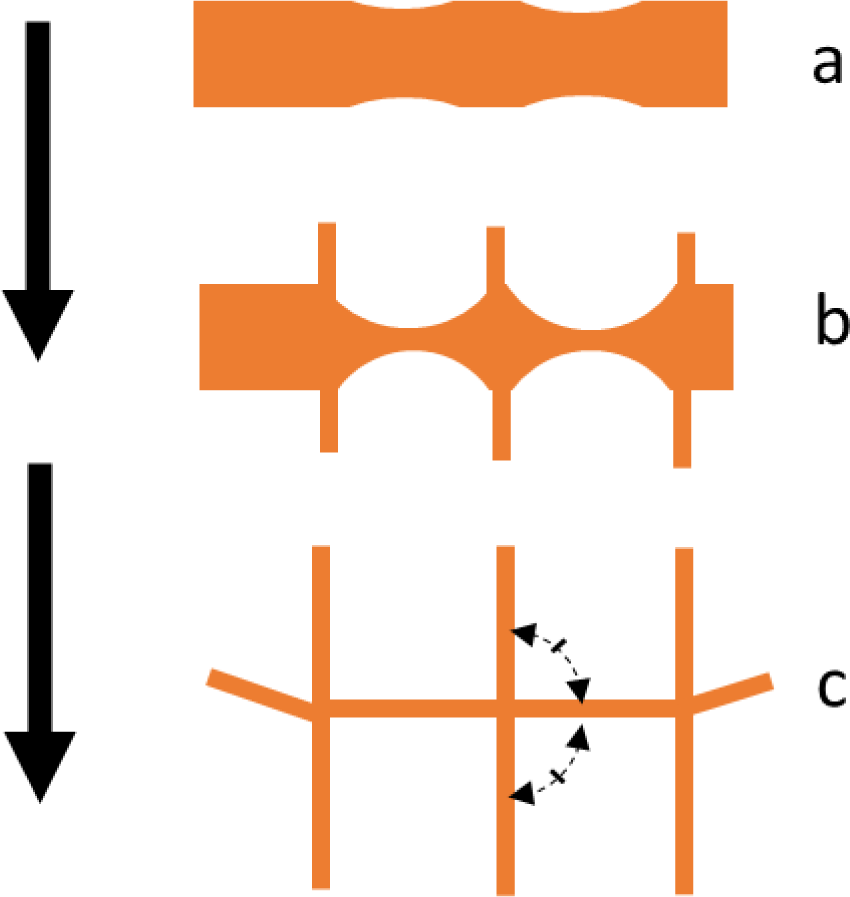
Predicted flat-bottomed cell production. (a) Cross section through the wax tab that carried the initial indentations to guide the alignment of cells. (b) The construction expected as initial cell formation by excavation of the guide pits and deposition of wax at their edges. (c) Further cell construction would thin the substrate and extend the cell walls. Cell location, guided by the indentations in the stimulus, would lead to cells aligned on opposite faces of the comb and, for those cell pairs, the 180 ° between opposing cell walls would be divided equally between the cells placing the mutual base orthogonal to them.

A thick wax backplane (∼1.2mm) was used to limit the interaction between work on both sides until the cell locations had been established and so reduce the tendency of the bees to build cells in a more natural alignment. Under conditions where cells on either side align, we predict that the mutual base will be built so the 180° between the side wall of one cell and that of its opposite will be shared equally causing the base to comprise a single facet at 90° to each cell side wall.

#### Experiment 3

This experiment was to test our third prediction that the initially uneven distribution of available space, both angular and area, will be redistributed between adjacent cells as construction progresses. Discrete tongues of comb, built independently, would likely be separated by a non-integer number of cell widths and have distinct orientations. At the time of contact between two such misaligned tongues the cells at the junction will exhibit a high degree of irregularity. If the prediction is correct, then following further construction, the cell sizes and corner angles would be redistributed and so irregularities in cell structure would be reduced from the initial levels by more than would be expected by chance.

Construction of several distinct tongues of comb was stimulated by preparing frames that carried four strips of wax approximately 25mm by 5mm and 0.5mm thick bonded, using molten beeswax, to the top bar of a hive frame (Fig. 1 d).

### Stimulus handling and construction time

Hive frames were prepared by bonding three stimuli tabs, using molten wax, to the underside of the top bar separated by approximately 20mm. At approximately 9 am, one such frame was placed into each honeybee hives and after approximately 4 hours it was removed to be inspected and photographed. Had insufficient comb construction occurred, then the frame was reinserted for a similar period. The stimuli bearing frames were photographed, initially and at each inspection, in natural daylight using a Samsung Galaxy Camera 2 EK-GC200. A jig was used to hold both the frame and camera at a distance of 390mm. The image, which covered slightly more than the while frame, had a resolution of 4608 by 2592 pixels. The frame, width of 333mm, covered approximately 3000 pixels resulting in an image resolution of approximately 9 pixels per mm. Image scale calibration was facilitated by the inclusion within the field of view of a graduated measure.

### Photographic record analysis

Alignment of photographs taken before and after a period of comb construction was performed using custom software written by the authors, FormImageCompare. This tool is also capable of accepting user input to mark the location of relevant features then using those points to compute their size, curvature or orientation. This software was written in C++ and developed using Microsoft Visual Studio Community 2019: Version 16.7.2, Visual C++ 2019. Image manipulation routines were supported by the library OpenCV:Version 3.3

The mechanical jig, used to support a frame and the camera, provided linear alignment within a few millimetres and angular alignment of a few degrees but this was insufficient for comparison of wax features at sub-cell scale. Alignment was improved using software to adjust one of each pair of images to make coincident each of four discrete points on a frame evident in both images allowing the position and shape of comb features to be compared.

#### Experiment 1 – Cells adjacent to a linear wax barrier

Cell construction against a straight wall stimulus was measured in three situations. Firstly, where adjacent cells (each of a depth <5 mm) had been built against one side of the stimulus. Secondly, where a wall found between adjacent cells was built against a barrier formed by the wooden frame. The third situation was where cell construction had progressed sufficiently for the stimulus to have been eroded and the cells had been built to a depth of several millimetres; a condition that was typically associated with comb having covered the test piece.

A sample cell’s side walls were measured from the photographs using the coordinates of points marked by the software operator, a point at each end of the sample cell wall, and further marks to locate the barrier (Fig. 4). Our custom software used the locations to calculate the orientation of each element and hence the angle between the wall and the barrier element.

**Fig. 4.**
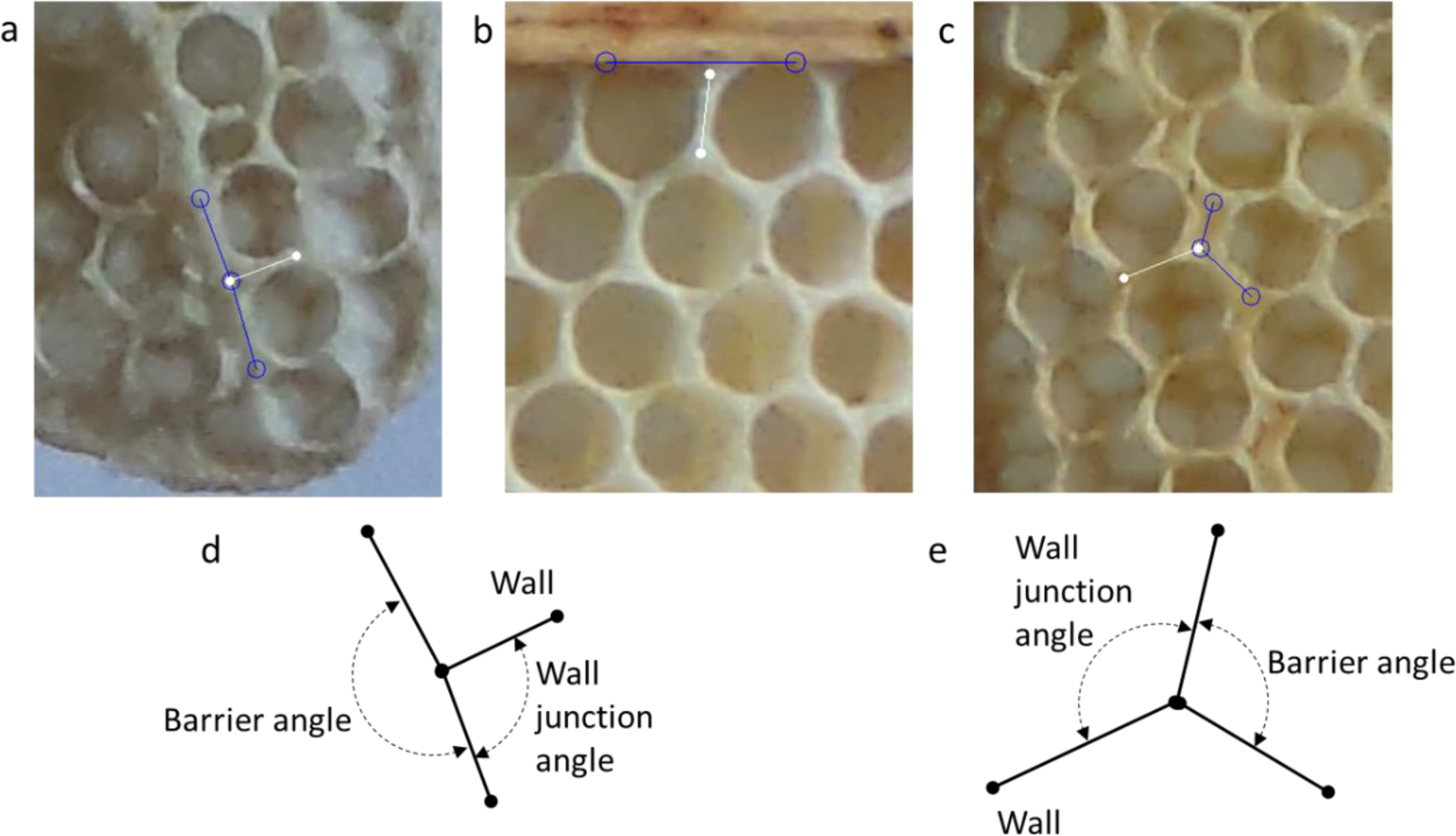
Example measurements taken of wall to wall angles. (a) A wall between a pair of cells built against a stimulus forming a wax barrier. (b) The wall between a pair of cells built against a fixed barrier, the wooden frame. (c): A later image of the cells shown in ‘a’ after further construction during which the workers had re-aligned the cell walls. (d) The measured attributes of the feature shown in ‘a’, the barrier angle and the wall to barrier angle. (e) Measurements taken after further construction, as shown in ‘c’, where the barrier had become eroded .

#### Experiment 2 – Opposing cells on either side of comb

Measurements were taken of cell bases found within comb formed on stimuli designed to encourage directly opposing cells (Fig. 1 b & c and Fig. 3). Bases selected for measurement were those within cells that were aligned base-to-base and showed a high degree of overlap, sufficient that when viewing the cell base via transmitted illumination no gap was perceptible between the side walls and those of the opposing cells (Fig. 5).

**Fig. 5.**
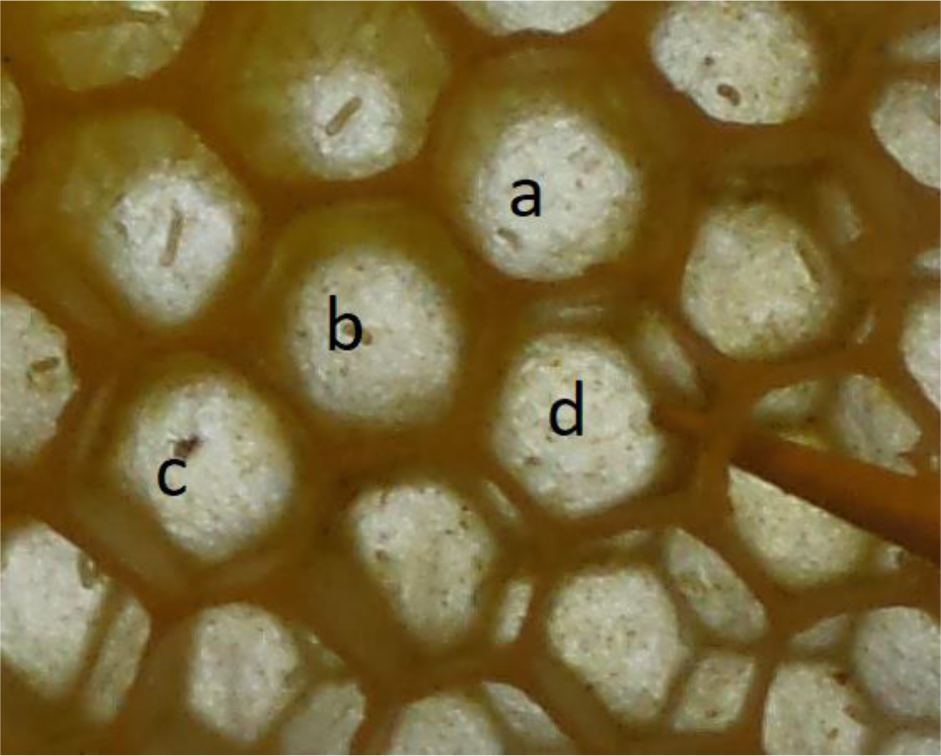
Flat-bottomed cell selection. The image shows, in transmitted illumination, an example collection of cells. Of these, the cells marked a, b and c were selected for measurements, but d and all others were rejected as they showed insufficient overlap.

Measurement of the bases was made using a multi-point depth gauge, which was placed into the cell to be measured, withdrawn and then photographed alongside a 10 mm gauge. Measurements were obtained from the photograph to provide the location of seven points (d0-d6) across the cell.

The control samples, used for comparison, were measurements of bases with cells found within natural comb the bees had added between experimental stimuli.

#### Experiment 3 – Cells at junction of merged sections of comb

For this experiment, measurements related to cell geometry were extracted from the photographs by out custom software. The initial state was where the tongues had made contact but had been separated in the immediately preceding photograph (taken 4 hours or less previously). Second-stage photographs were of the comb following between eight and twelve hours further activity, recording the post-merger condition of the cells.

Image processing involved alteration of brightness and contrast, followed by conversion to black and white using a localised threshold value computed using a Gaussian filter over an area of 63x63 pixels, which for these images was approximately the size of a cell. Region boundaries computed for the resulting binary image were then filtered to leave those with an appropriate area and proximity to other such zones. For each zone an external hull, a shape comprising only convex external vertices, was used to derive the location of its centroid, then used as the centre of a cell.

This image processing, and drawing of the resulting cell mesh, was performed by our custom software in real-time, thus the operator could manipulate on-screen controls to adjust brightness, contrast, threshold and other values allowing the user to interactively optimise the outcome (Fig. 6 - c).

**Fig. 6.**
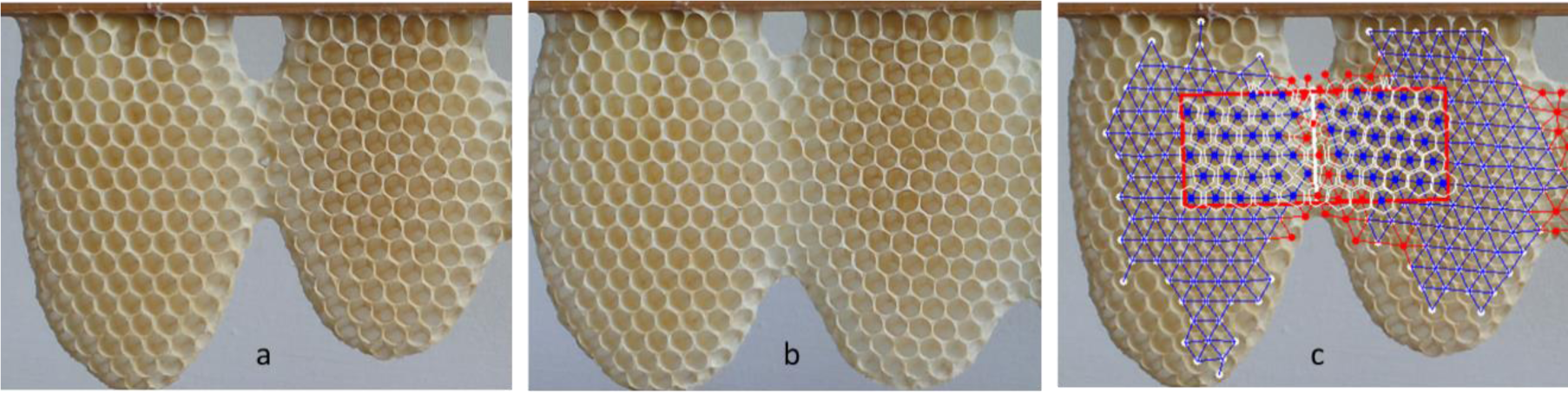
Merged tongues of the comb. (a) Tongues recently merged and (b) the same comb with an additional 10 hours of construction. (c) The software identified cell structure including the area marked for analysis.

Processing each image yielded coordinates for more than 95% of the cells and the remainder were then explicitly identified by the operator.

Using the centre of each cell, the software computed the distance to each adjacent cell. It also estimated the location of a mutual wall as the mean of the centre to centre midpoint and the point of maximum brightness on that line. The remaining aspects of cell geometry, number of walls, wall lengths and internal angles, were derived from the centroid and wall intersections.

Using the earlier image of a pair, showing the initial contact between the tongues, the operator marked each end of a tongue merger junction. The software used these two locations to define the centre line of a zone of interest and, using these, computed a rectangle measuring 30 mm on either side of the centre line. Cells with centres contained within that rectangle were those sampled.

The software used the location of cells to match those in one image with the cells in the second image allowing analysis of cell by cell alterations in their geometry.

### Analysis

Data obtained from the images, using FormImageCompare, was subsequently processed using custom scripts written in R, executed on R runtime version 3.6.3 within RStudio version 1.3.1093.

#### Experiment 1 – Cells adjacent to a linear wax barrier

Measurements were made of the angle between a cell wall and a constraining barrier and the value was compared with the predicted value of 90°. Where the barrier was eventually eroded, giving rise to a three-way junction, each of the three angles were compared with a predicted value of 120°.

A Two One-Sided Test (TOST), implemented in R by the function TOSTone(), was used to compare the measured angles with the predicted value (either 90° or 120°). For these tests we used the Smallest Effect Size of Interest (SESOI) of 0.5 standardised mean difference, Cohen’s D.

#### Experiment 2 – Opposing cells on either side of comb

Measurements taken of the bases using the multi-point depth gauge yielded the depth at seven points. The maximum displacement from the regression line through all seven points on the profile (d0-d6) was used as the measurement of flatness.

The base angle, one each side of the cell between the base and the vertical side of the cell, was calculated by taking the gradient of a regression line through each half of the base profile, from d0 and from d6 to the deepest location, typically d3 or d4.

A TOST comparison between the measured base inclination and the predicted value of 90° was performed with the SESOI of 0.5 standardised mean difference.

#### Experiment 3 – Cells at junction of merged sections of comb

The measurements gathered for each cell, its geometry and position relative to the cells that surround it, are not independent of those for adjacent cells: for example where two cells share a wall. Artificial duplication was avoided by using only data that applied to a feature outward of the cell centre; away from the junction.

The cells exhibiting the greatest irregularity were expected to be those close to the line of contact between tongues, therefore cells less than two cell’s width from the junction (10mm) where used for comparison between stages 1 and 2.

The metrics used for comparison were found to not be normally distributed and therefore a Wilcoxon test of comparison between values at each stage was performed using the R function wilcox.test(). Our prediction was that continued construction would optimise the cell layout causing a change between stage 1 and stage 2 reducing the irregularities in cell structure more than expected by chance.

The features measured to asses comb regularity were separation, distances between a cell’s centre and those of neighbouring cells, a cells internal corner angle, its wall lengths and its area. For each of cell separation, internal angles and wall lengths the metric of regularity was calculated as the standard deviation of those values.

Cells for which the location could be matched between stage 1 and stage 2 allowed the analysis of changes made to individual cells. Cells were deemed to match where the location differed by less than 2.5 mm.

## Results

### Experiment 1 – Available angular range was divided between the cojoined cells

Measurements were made of 73 cell walls connected to a stimulus barrier and shared by two adjacent cells (Fig. 7). The corner angles were measured as 90.9°± 7.0° (means ± standard deviation, throughout). Measurements were also taken of 72 cells at the top row of comb built against the frame for which the corner angles measured at 90.0° ±

**Fig. 7.**
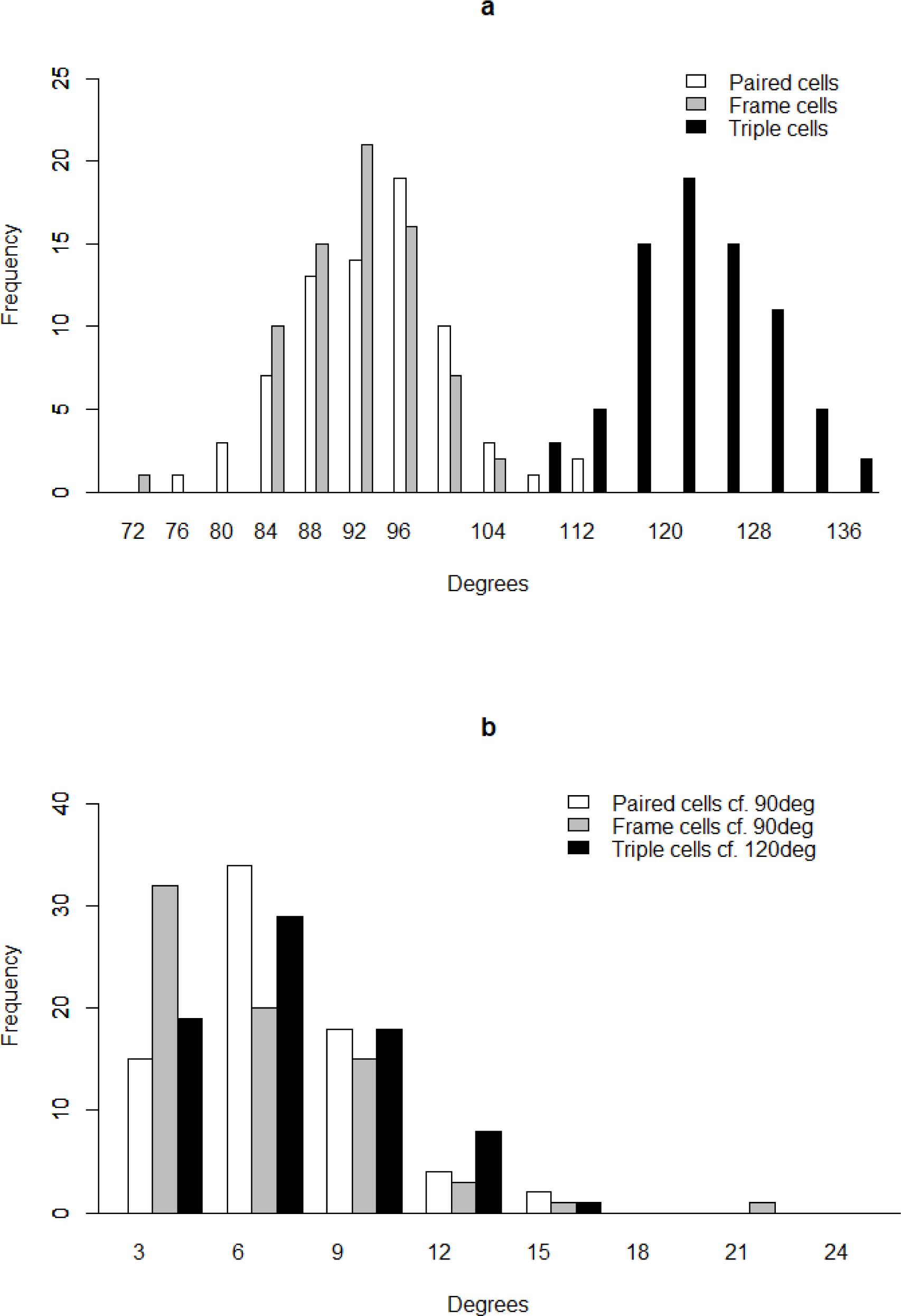
Cell corner angle measurements. (a) Distribution of the angle at cell corners in three configurations. Paired cells are those at a junction with the barrier where the wall was between two cells (N = 73 and mean = 90.95°, predicted to be 90°). Those labelled ‘Frame cells’ are walls shared by cells abutting the wooden frame, (N = 72 and mean = 89.96°, predicted to be 90°). “Triple cell” measurements were taken from the 3-way wall junctions where the barrier had been eroded (N = 75 and mean = 119.40°, predicted to be 120°). (b) The deviation of those cell corner angles from the predicted angle, for each condition, means of 5.24°, 4.21° and 5.29°.

5.53°. A sample of 75 cells that were initially built against a barrier but subsequent construction had eroded the barrier were also measured. These corner angles were measured as 119.4° ± 6.46°.

Samples for each junction type, paired, triple and frame, were compared for equivalence to the predicted values of 90°, 120° and 90° respectively. The equivalence of the paired samples and the frame samples was also tested. For paired cells against a linear wax barrier, the corner angles were equivalent to 90° (P=0.0013 & 0.249; probability of different from reference/probability of equivalent to reference) and triple cell corners were equivalent to 120° (P=0.00036 & 0.42). For paired cell corners against the frame were equivalent to 90° (P=0.00004 & 0.955), and paired cells corners were equivalent to frame cell corners (P=0.020 & 0.346).

These results demonstrate that the angles between cell walls are balanced to equalise distribution of the available angular range, as predicted.

### Experiment 2 – Single facet base was built orthogonally to the walls of two opposing cells

Measurements of the bases shared by the experimental opposing cells and of further cells sampled from natural comb. The contrasting shape of the bases found within these sample set is illustrated (Fig. 8) showing the height for each probe point relative to the ends. The heights are presented as boxplots showing median, interquartile range and outliers.

**Fig. 8.**
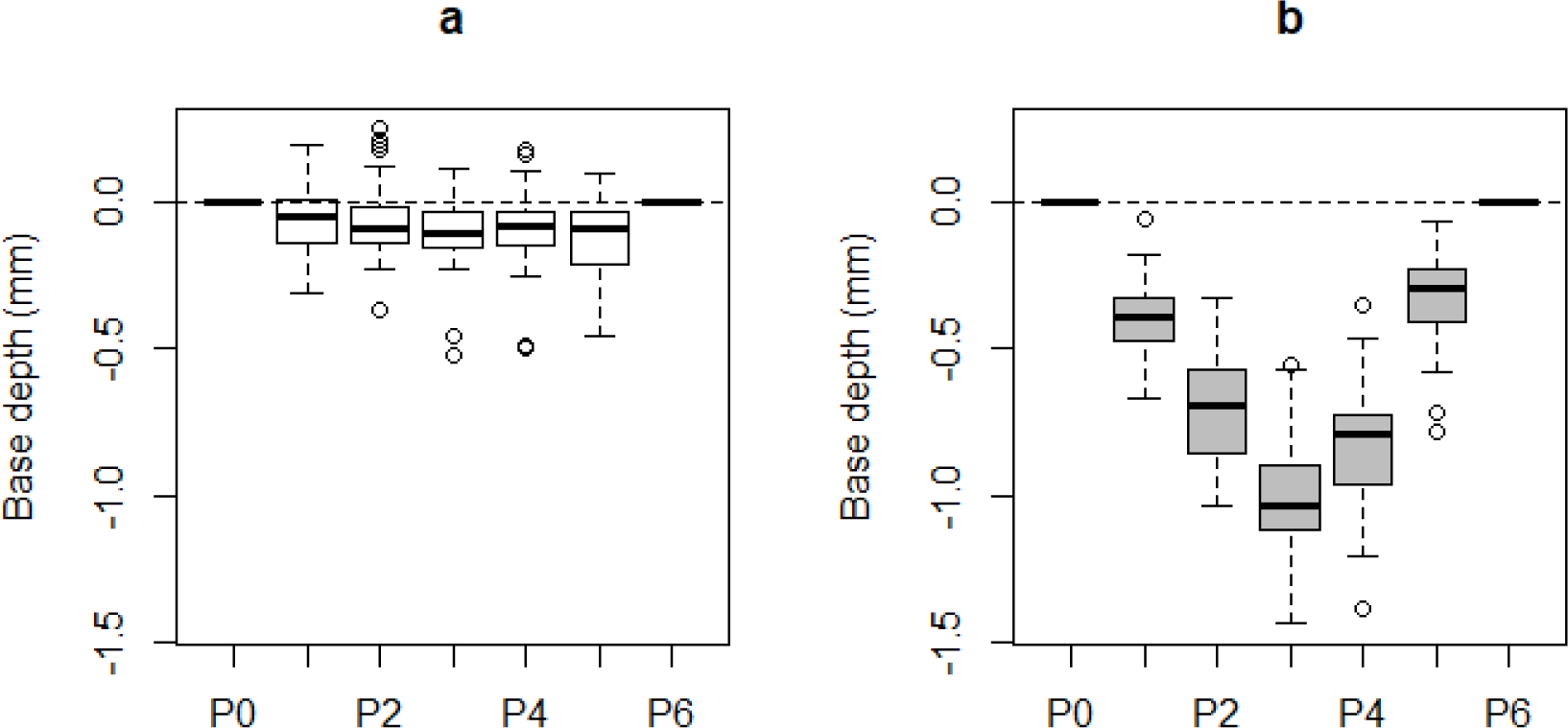
Profiles comparing the bases of opposing with non-opposing cells. (a) Profiles for bases where a cell on one face is directly opposite a cell on the other face. (b) Profiles of bases within cells that formed naturally and thus a cell on one face was typically offset by half a cell width from cells on the opposing face. Bases in opposing cells (a) comprised a single, flat, face at right angles to the cell-walls, whereas natural cells (b) have pyramidal bases with a ‘V’ shaped profile..

Measurements were made of the bases of 42 cells, each in opposition with a cell on the other face of the comb. The maximum distance between the base and the line of regression through the 7 points on the profile was 0.13 ± 0.06 mm, which was significantly less than that measured for cells within natural comb (0.58 ± 0.11 mm; t2=65 = 22.59, P <0.00001; Fig. 9).

**Fig. 9.**
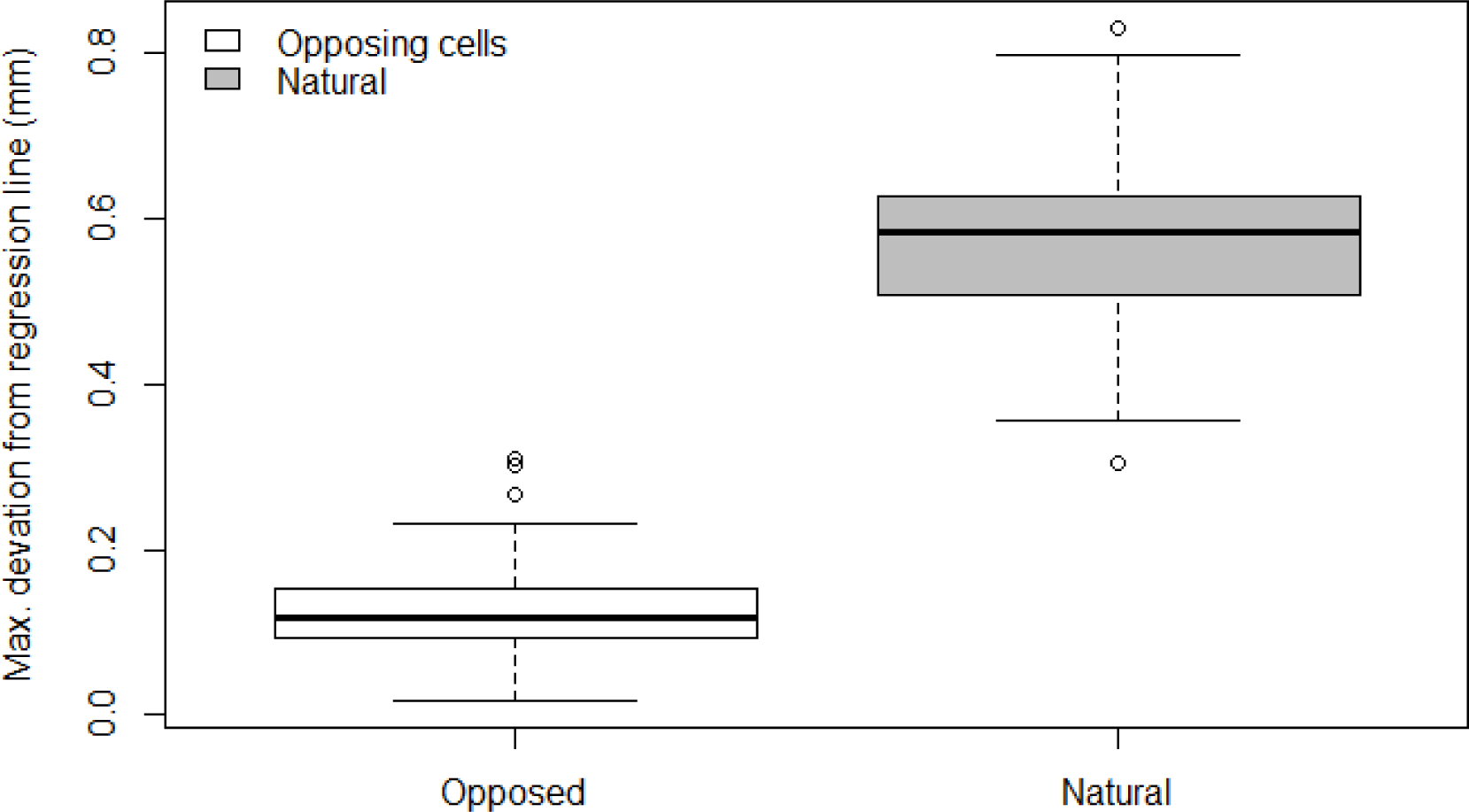
Maximum deviation from a regression line through the seven depths acquired with the multi-point probe, used as a measurement of flatness. The results shown are for opposing cell bases (N=42, mean=0.13) and for the bases of natural cells (N=43, mean=0.58).

The slope of the base relative to the side walls of its cell for those in opposition was measured as 90.3°± 6.7°, which was significantly less than that for each side of a base from a natural cell (116.2°± 7.5°; t65 = 22.59, P <0.00001; Fig. 10).

**Fig. 10.**
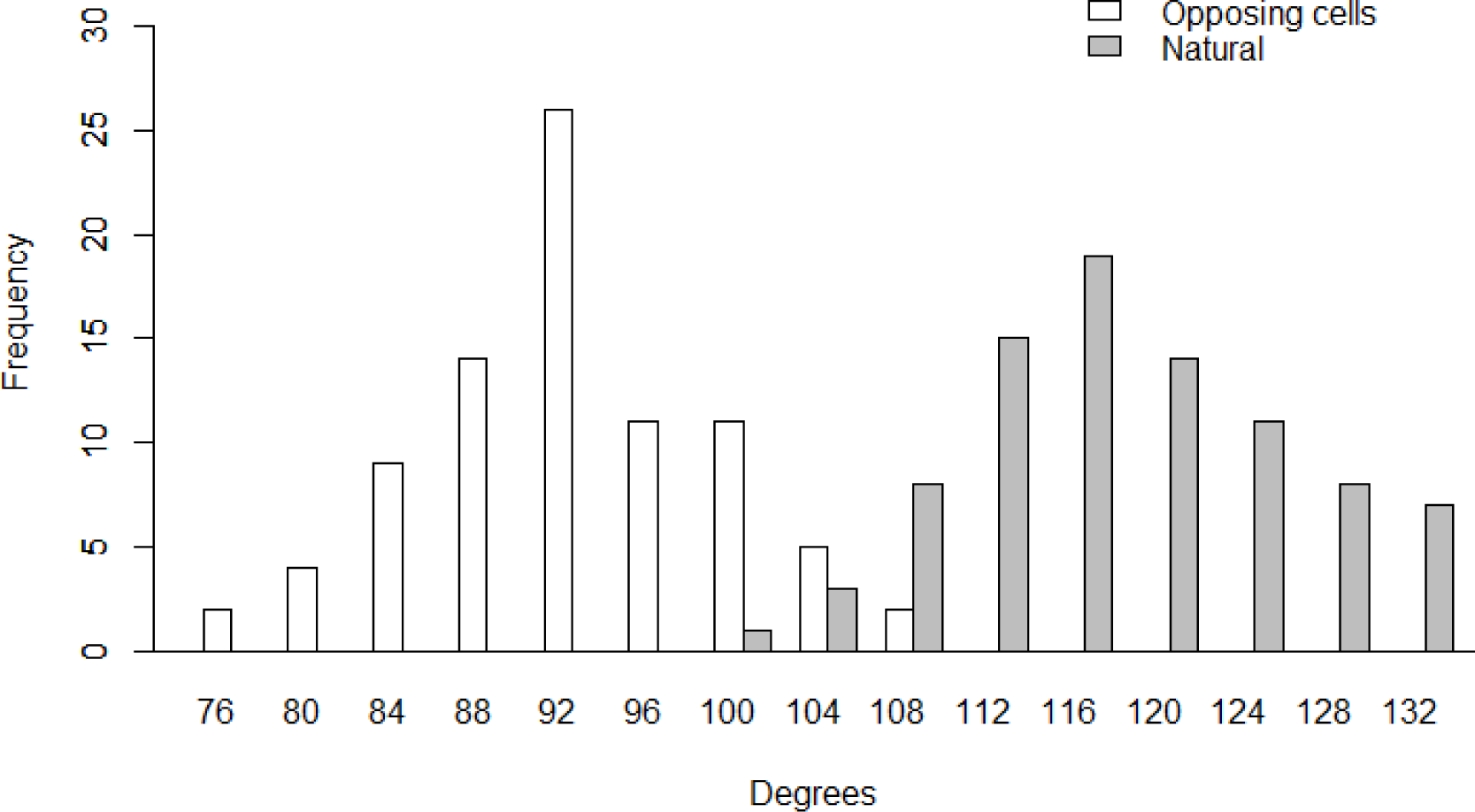
The distribution of angles between the base of a cell and its side wall. The samples of bases in cells in opposition (n=84, 90.3°± 6.7°) are distributed distinctly from those for natural cells (n=86, 116.2°± 7.5°).

The angle of bases to the side wall of cells in opposition was found to be equivalent to the predicted value of 90° (TOST with SOSEI of 0.5 standardised mean difference, Cohen’s d; P=0.000048/0.673; probability of difference from reference/probability of equivalent to reference).

### Experiment 3 – Cell layout was optimised after discrete sections of comb were connected

Measurements were made of cells within 30mm of the line of contact where two tongues touched, of which 1315 were measured at stage 1, shortly after the contact, and 1272 were measured at stage 2, following further construction. Of these cells 458 and 429 respectively were close to the line of contact (within 10mm). The geometric characteristics of cells used for these comparisons were inter-cell separation, intra-cell corner angle, wall length and cell area.

All measurements of irregularity were greater for cells closer to the junction. For cells within 10 mm from the junction the features of cell geometry were compared between stages 1 and 2.

#### Cell separation

The standard deviation of centre to centre separation of a cell from its neighbours at stage 1 was 0.52 ± 0.26 mm, which was significantly greater than that at stage 2 (0.37 ± 0.21 mm; P <0.00001; Fig. 11). This demonstrated that cell wall positions were altered to balance the space allocation between adjacent cells, as predicted.

**Fig. 11.**
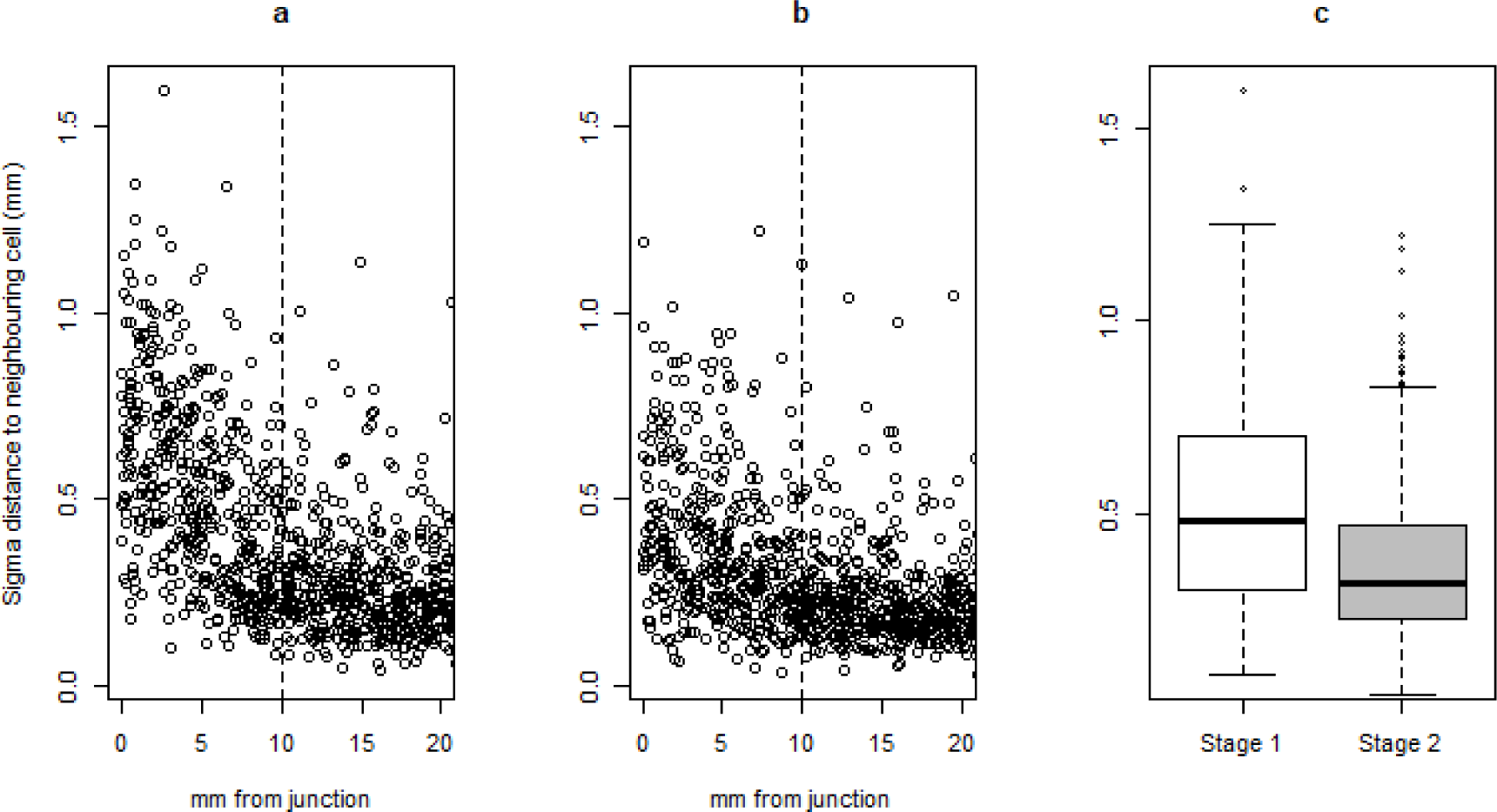
Distribution of inter-cell distance. (a) Standard deviation of the distance between a cell and its neighbours at stage 1, for each cell, plotted against the distance of the cell from the junction and (b) the same metrics but measured after further construction, at stage 2. (c) Measurements of those cells within 10mm of the junction (points on a & b to the left of the dotted line) at stage 1 and at stage 2 showing the comb had became significantly less irregular.

#### Corner angle deviation

The standard deviation of corner angles around a cell at stage 1 was 0.15 ± 0.09 rad, which was significantly greater than that at stage 2 (0.10 ± 0.06 rad; P <0.00001; *Fig. 12*) showing that cell walls were repositioned to even their layout as predicted.

**Fig. 12.**
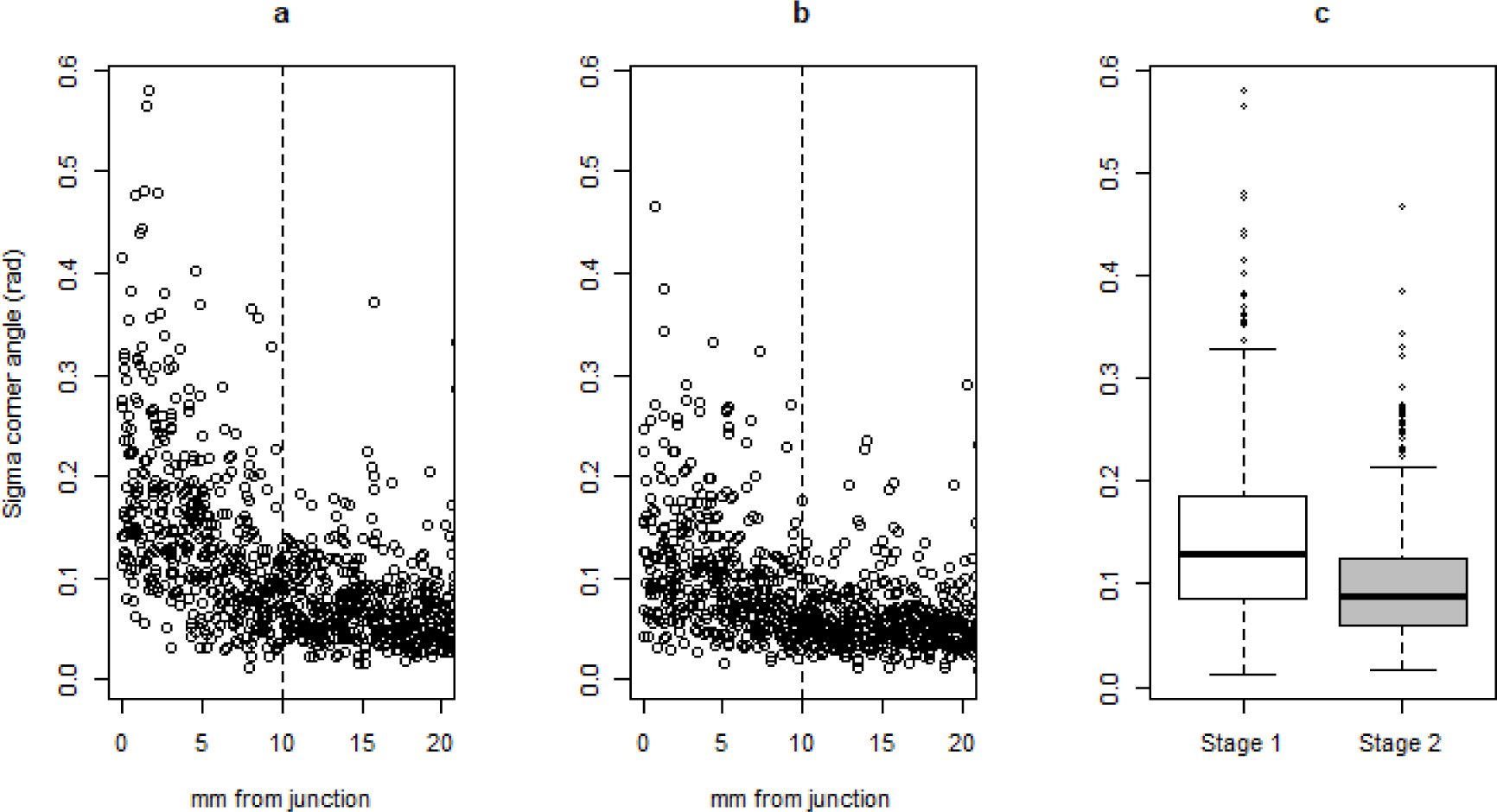
Distribution of cell internal corner angles. (a) Standard deviation of the cell corner angles at stage 1 plotted against the distance between the cell and the junction and (b) the same metric measured at stage 2. (c) Sampling those cells within 10mm of the junction (points to the left of the dotted line), the measurements taken at stage 1 and those taken at stage 2 showing the comb had become significantly less irregular.

#### Wall length

The standard deviation of the length of walls around a cell at stage 1 was 0.72 ± 0.35 mm, which was significantly greater than that at stage 2 (0.55 ± 0.26 mm; P <0.00001; Fig. 13). This demonstrated that cell walls were repositioned to even the perimeter, as predicted.

**Fig. 13.**
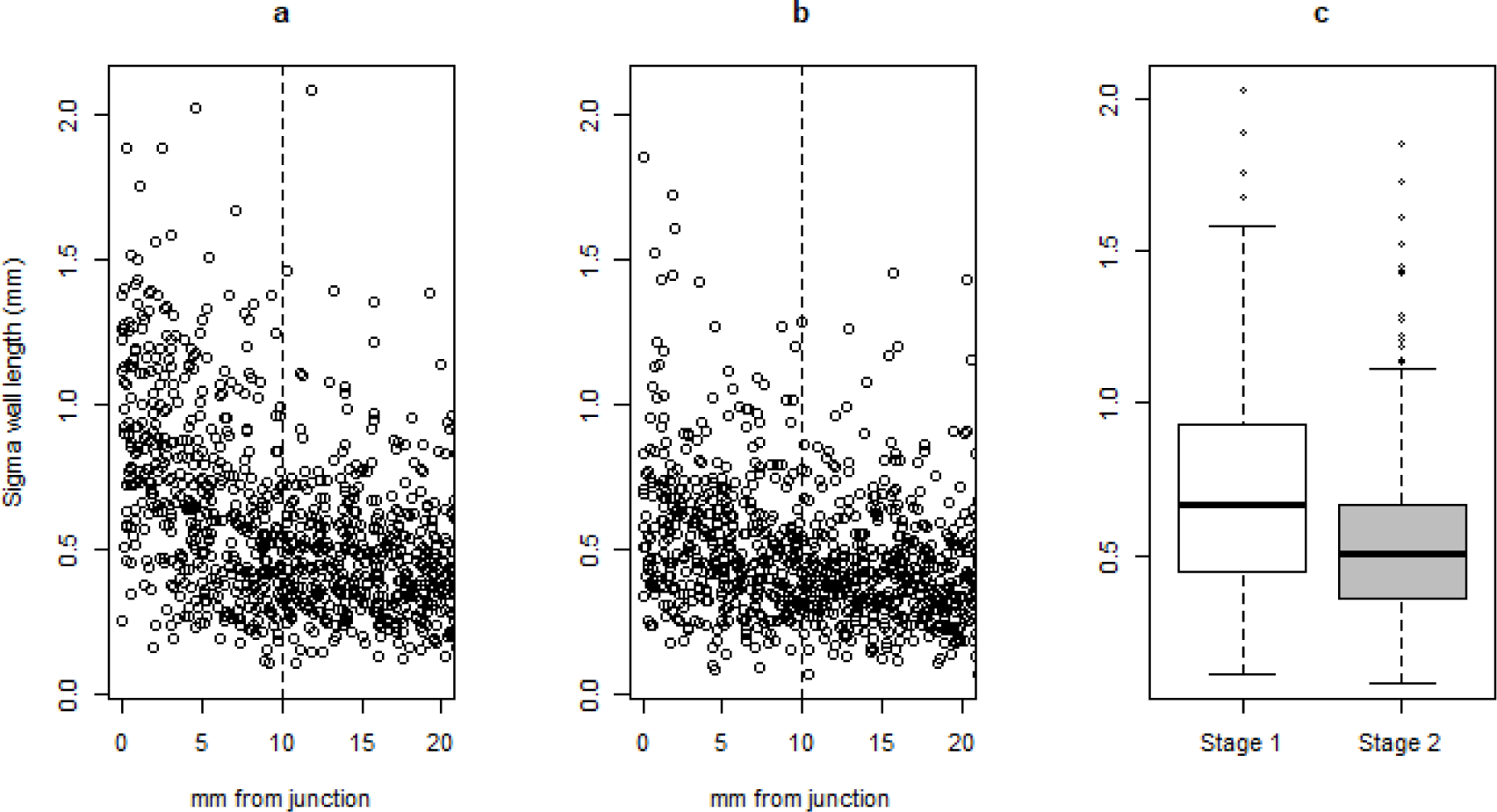
Distribution of cell wall lengths. (a) Standard deviation of the lengths of the walls around each cell at stage 1 plotted against the distance between the cell and the junction and (b) the same metrics but measured at stage 2. (c) Sampling those cells within 10mm of the junction (points to the left of the dotted line), the measurements taken at stage 1 and those taken at stage 2 showing the comb had become significantly less irregular

#### Cell area

The area of cells close to junctions (< 10 mm) was compared with a reference cell area, that being the mean of areas of cells further away from the junction (>20 mm). The reference area for more distant cells at stage 1 was 24.19 ± 4.0 mm^2^ (n = 431), and 24.16 ± 3.0 mm^2^ (n= =413) at stage 2. The reference area was thus computed as the balanced mean area of cells from both stages, 24.17 mm^2^. The cells sampled at stage 1 were undersized compared to the reference area at stage 1 by 3.38 mm^2^ ± 5.0 mm^2^, which was significantly greater than that at stage 2 (1.31 mm^2^ ± 4.1 mm^2^; P <0.00001; Fig. 14). This demonstrates that cell walls were enlarged, closer to the mean size, as predicted.

**Fig. 14.**
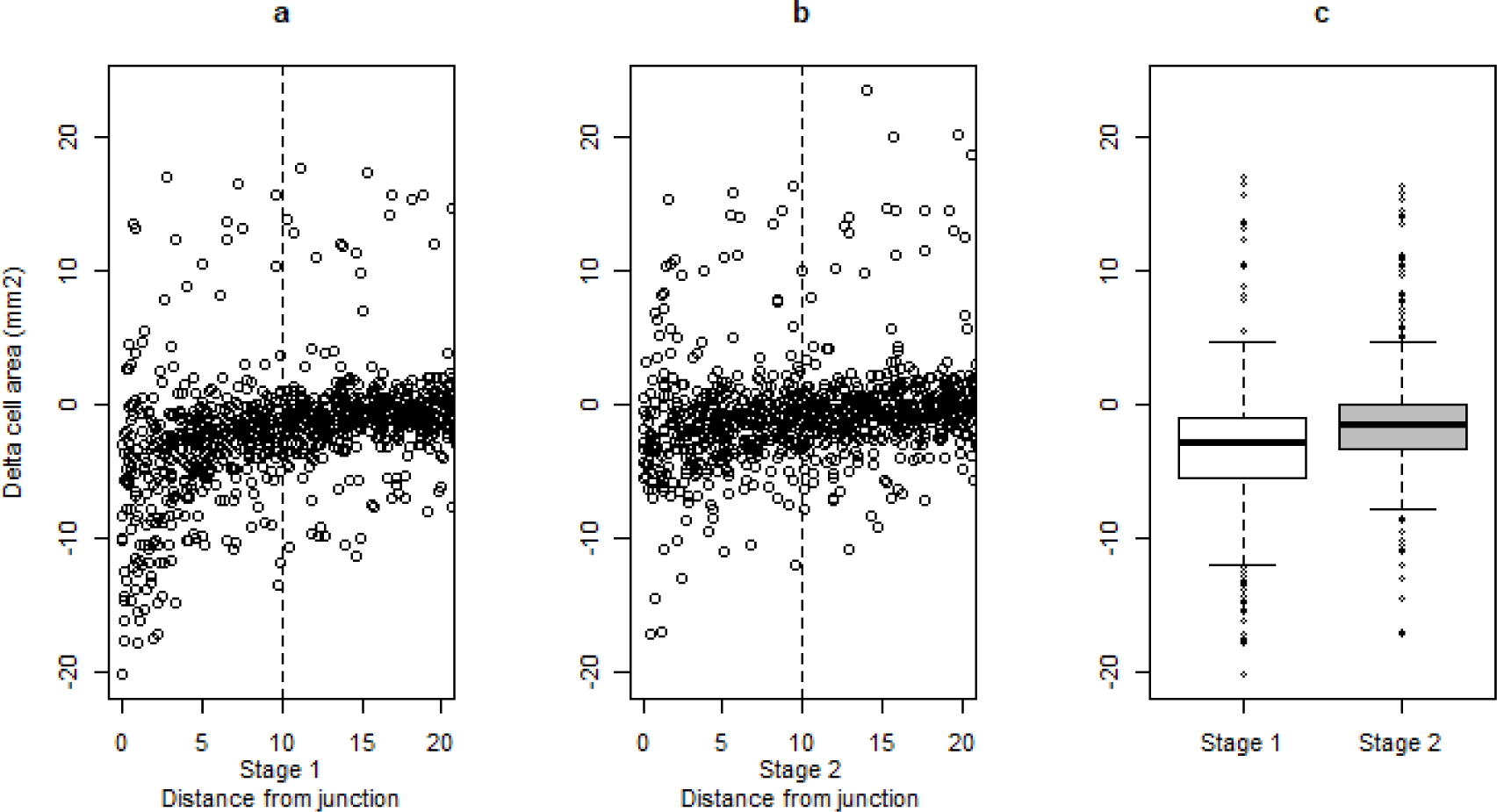
Distribution of cell area as a function of distance from the tongue junction. (a) The difference between the area of a cell and the reference cell area at stage 1 plotted against the distance between the cell and the junction and (b) the same metric but measured at stage 2. (c) Sampling those cells within 10mm of the junction (points to the left of the dotted lines in a & b), the measurements taken at stage 1 and those taken at stage 2 show that the cell area had increased significantly, towards the reference mean, as the comb layout had become more regular.

#### Cell-by-cell comparison

The results presented above show the differences between the whole of the sample sets at stage 1 and stage 2. By matching cells between images, we were able to identify the change in each cell from stage 1 to stage 2. For cells close to the junction (< 10mm) we paired 409 of the 458 cells (89.3%) at stage 1 with 409 of the 429 cells (95.3%) at stage 2.

#### Cell-by-cell comparison of corner angle deviation

Of the 409 cells within 10mm of the junction, variation in corner angles was reduced between stage 1 and stage 2 in 282 cells (69%) and we found a significant change towards the sample mean at stage 2 (t(408) = 10.53, P < 0.00001), with 288 cells (70%) altered to be closer to the mean (Fig. 15).

**Fig. 15.**
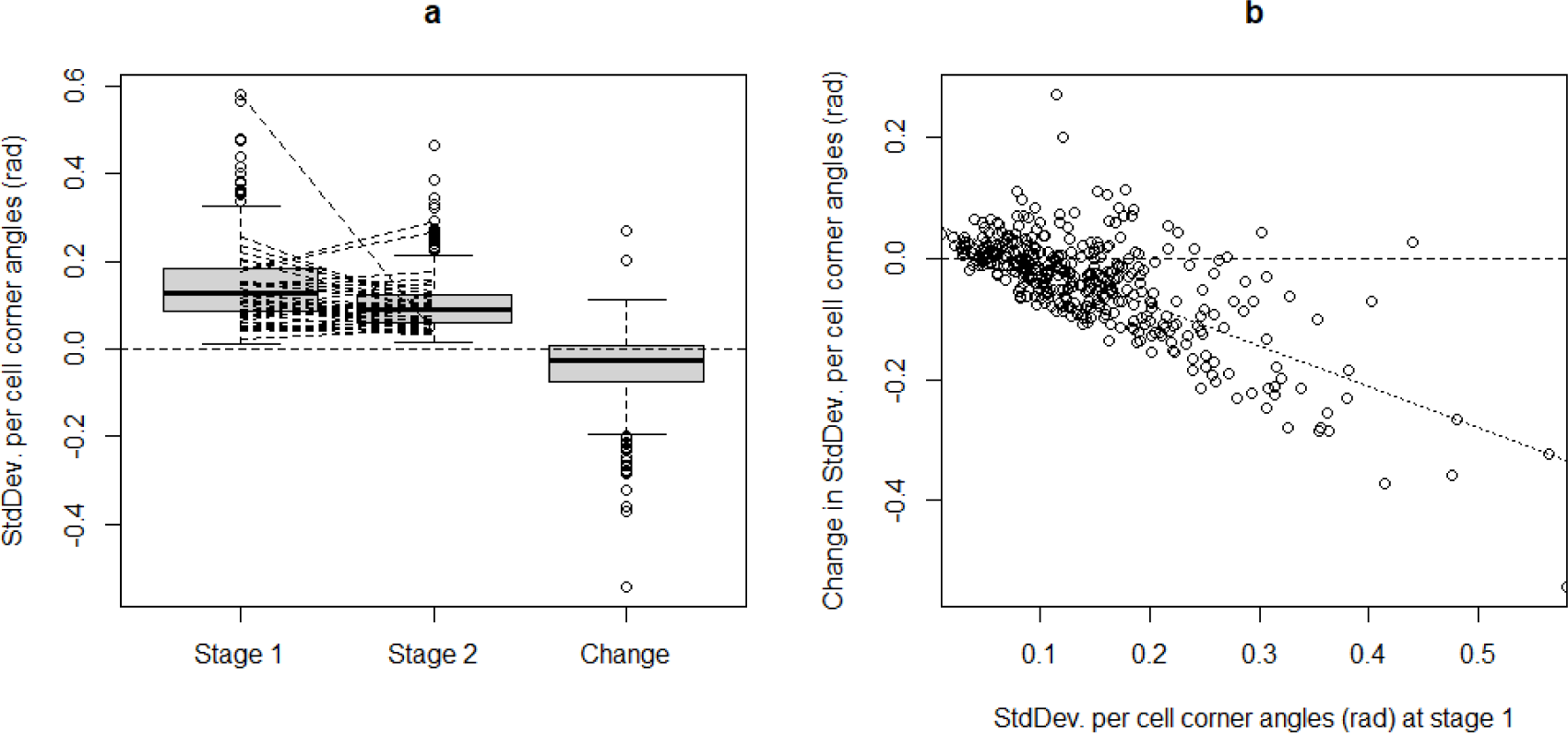
Reduction of angular irregularity. (a) Distribution of cell corner angle irregularity at stages 1 and 2. The solid line on each plot shows the median value, while the box shows the interquartile range for the samples. Additional lines from stages 1 to 2 show the individual cell changes. For clarity, only 10% of the changes are shown. The distribution of change for each cell is also shown (282 out of 409 showed reduced irregularity). (b) The change of cell corner angle irregularity is plotted against that measure at stage . These values are related, having a Spearman rank correlation of - 0.62. The points are overlayed by a regression line relating the values and indicating that the more irregular cells underwent greater improvement (ratio of change to original value = -0.67).

The improved regularity of cell corner angles can be seen in Fig. 15 b. The change in cell corner irregularity was found to be correlated with the difference between that value at stage 1 and the mean at stage 2 (Spearman’s rank-order correlation ρ(407) = -0.62, p<0.0001) showing that the more irregular cells were changed toward that mean by the greater amount.

#### Cell-by-cell comparison of cell area

Of the 409 paired cells, 275 (67%) showed an increase in cell area between stage 1 and stage 2. A significant change towards the sample mean at stage 2 was identified (t(408) = 10.53, P < 0.00001), with 289 cells (71%) found to be closer (*Fig. 16*). The improved regularity of cell area that occurred between stage 1 and stage 2 can be seen in *Fig. 16* b. The change in area was found to be correlated with the difference between that value at stage 1 and the mean at stage 2 (Spearman’s rank-order correlation ρ(407) = -0.54, p<0.0001) showing that cells with an area that differed from the eventual mean by the greater amount (either larger or smaller) were changed most towards the eventual mean.

**Fig. 16.**
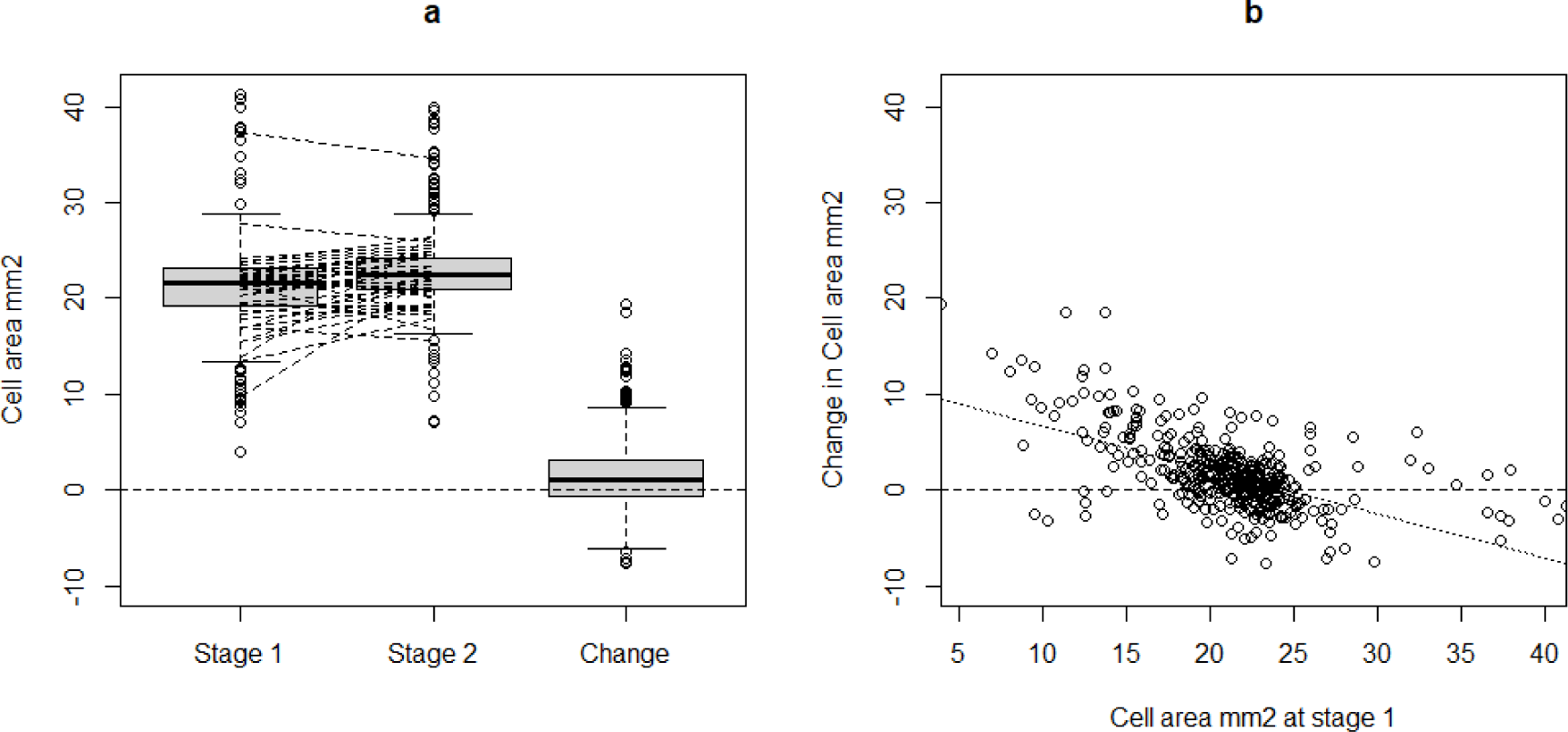
Improved consistency of cell area. (a) Distribution of cell area at stages 1 and 2. The solid line on each plot shows the median value, while the box shows the interquartile range for the samples. Additional lines from stages 1 to 2 show the individual cell changes. For clarity, only 10% of the changes are shown. The distribution of change for each cell is also shown where 275 out of 409 cells increased in area and 289 out of 409 changed toward the stage 2 mean. (b) Change in cell area is plotted against the area at stage 1, values that are related having a Spearman rank correlation of -0.54 The points are overlayed by a regression line(dotted) relating the values and indicating that undersized cells were enlarged while oversized ones were made smaller (ratio of change to original area = -0.46).

### Discussion

Honeycomb has a pleasingly regular form, comprised of almost flat blades formed from nearly regular hexagons. The regular sized hexagons lay in clear, nearly straight, rows of cells that combine to form a structure that has the appearance of being strictly defined and intentional. Many writers have commented on and discussed the appearance and structure of honeycomb with perhaps the earliest analysis of being attributed (Heath 1921) to Pappus of Alexandria who observed that honeybees make “honeycombs,(with cells) all equal, similar and contiguous to one another… they have contrived this by virtue of a certain geometrical forethought. … the figures must be such as to be contiguous …Bees, then, know just that the hexagon is greater than the square and the triangle and will hold more honey for the same expenditure of material used in constructing the different figures.” Closer inspection of comb reveals that these aspects are only approximately regular (Hepburn and Whiffler 1991). Each cell, and indeed each wall of a cell, is formed subject to the builders’ ability to respond to local features, which invariably leads to local inaccuracies (Smith et al. 2021).

Kepler (1611) wrote of the bees’ choice of a hexagon, with its internal angles of 120°, is the polygon best suited to the tessellation of cells across the plane of honeycomb. Kepler also explored the geometry of honeycomb cells bases, his analysis of which resulted in the first description of a rhombic dodecahedron, a regular solid with internal angles of 120° and one that therefore tessellates in three dimensions. When considering a potential mechanism that gave rise to the hexagonal outline, and the pyramidal bases, of cells, Kepler (1611) compare comb with other naturally occurring semi-rigid objects deformed in a geometrically regular fashion. His first example was of peas in a pod. The pod linearly constrains the peas with the result that they each adopt a flat face that is orthogonal to the line of constraint. This example from Kepler compares well with the outcome of our experiment 1, where the angular range was restricted to 180° and when distributed between two cells resulted in corner angles of 90°.

The base of a cell, comprising three rhombi each shared with a cell on the opposite face, is shaped as a pyramid which could be of any depth but the geometry of the base determines both the depth of the pyramid and the angles within the rhombi. The internal angle of the rhombi where they meet at the centre of the cell could be slight less than 120° for a nearly flat base and increased base depth would cause that angle to reduce, approaching zero for extremely pointed base. The major angle of a base rhombus was measured to be 109° 28’ (Maraldi 1712), a value cited by Huber (1814, p. 107), but additionally Huber considers whether this base geometry is the most efficient and, citing Réaumur and Koenig who calculated the ideal angle to be 109° 26’, Huber and others considered that the bees built comb in order to maximise the storage volume for a minimum quantity of wax. An alternative mechanism that leads to the observed geometry is provided by the outcome of our experiment 1 for triplets of cells positioned within an angular range of 360° that results in an even distribution causing each cell to occupy 120°. When applied to three cells in a 2-dimensional arrangement, the result is a triple junction within the tessellated hexagons visible on the face of the comb. However, when the junctions are equalised at the cell base between four cells in a 3-dimensional arrangement, the results from experiment 1 show that each corner will settle at dihedral 120°. The base will therefore adopt the shape of a regular rhombic dodecahedron within which each dihedral angle is 120°. The resulting rhombi angles can be calculated to be 2 * Cos^-1^(1/√3) or 109° 28.27’. This implies that bees do not construct comb in a manner aimed at maximising efficiency, rather, the more parsimonious explanation is that they simply balance the distribution of angular space between adjacent cells.

The discussions above concern the depth of the basal pyramid being either exactly, or close to, a section of a rhombic dodecahedron, but we saw in experiment 2 that bees built abnormal cells with flat bases. While such cells differ from those normally found in naturally produced com, this unusual layout was nonetheless predictable. Our prediction that the cell internal corner angle will be an integer fraction of the total range does not state that every angle within a cell should be 120°. For a range of 180° and where the number of cells is two, then the wall angle would be predicted to be 90° – the outcome of experiment 2. Flat bases were formed where cells lay in direct opposition, while normal rhombic bases were formed where cells overlapped by half their width, with the centre of a cell on one face positioned opposite the triple junction on the other face. Our prediction addresses both situations, but where the cells are positioned in different arrangements, the same mechanism will form alternative base structures, hence the geometry and arrangement of base faces arises as a consequence of the alignment of cells on either side of the comb. Fig. 17 shows a range of base forms including pyramidal bases, flat bases and some that approximate the truncated octahedron which Tóth (1964) suggested offered improved efficiency. In all cases, each facet of the base, a flat side, would be formed where a cell on one face overlaps one on the opposite side of the comb. A similar range of base styles was constructed when naïve bees built comb without prior experience in doing so (Oelsen and Rademacher 1979). The comb built by these bees lacked rotational alignment between cells on each face, and this resulted in a circular pattern of cell overlap, which was described as ‘floral’ by the authors.

**Fig. 17.**
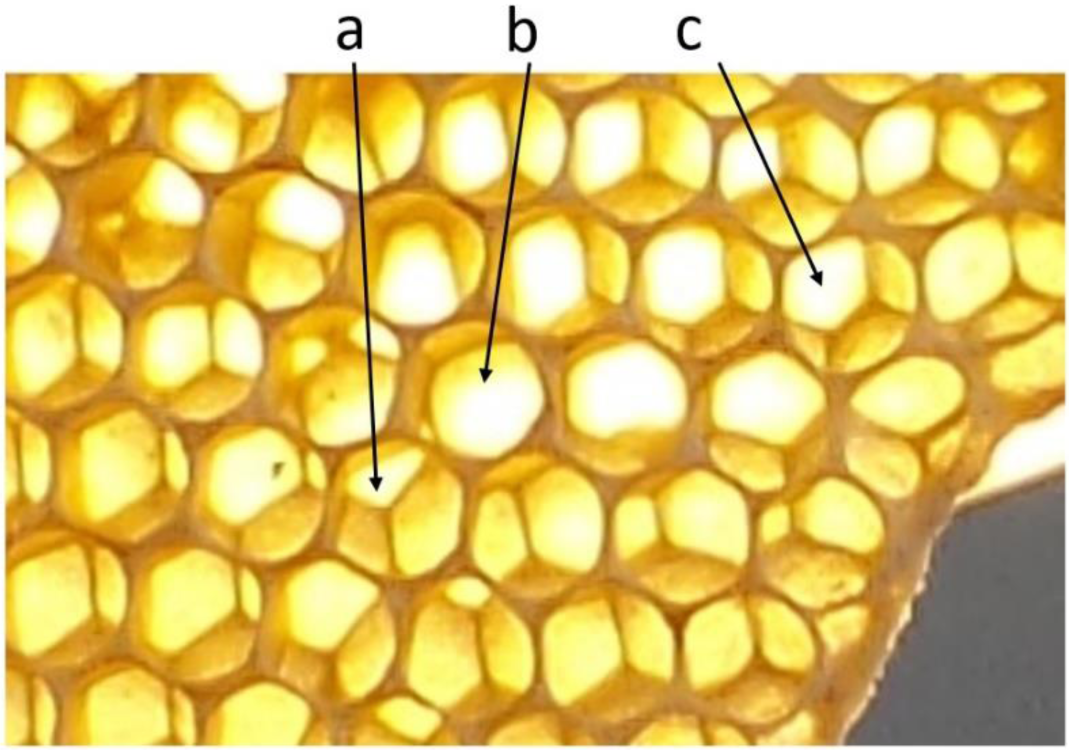
Varied base forms resulting from cell alignment found in a section of natural comb. Cell ‘a’ is aligned with those opposite in a near-conventional fashion so has a rhombic base. The cells either side of the base ‘b’ are directly opposite each other causing a flat base and the centre of cell ‘c’ is aligned above the side wall of opposite cells and the resulting base is a section of a truncated octahedron. The pattern of cell overlap around ‘b’ is similar to that described as ‘floral’.

Cell adjustment has been observed, but not explained, where bees were offered wax foundations on which to build comb where the indentations included those of an inappropriate scale (Yang et al. 2010). The authors found that the larger bees, *A. mellifera*, built cells on foundations with indentations that were spaced too closely (being appropriate for *A. cerana*) but they did so by extending the initial guide. An explanation for this is suggested by the outcome from for our experiment 3, demonstrating a progressive adjustment of each cell towards the appropriate size. The location of cell walls would initially be guided by the foundation but, as the authors observed, cell walls were subsequently relocated to form cells of the correct size for specific workers, albeit with a biproduct of malformed cells and at the expense of comb scale regularity.

Coercing bees to build comb on inappropriate foundations creates the conditions for comb construction to start irregularly over a wide area but irregular cells are also created along the line of contact when two tongues of comb merge, although ‘collide’ may be the mot juste. Cell level irregularities in wall length and cell area at and around intersections were found to change smoothly across a distance of four or five cell widths to either side of a junction (Smith et al. 2021). The authors posited that the smooth progression may result from the bees anticipating the narrowing gap and, using touch to detect the relative alignment, determine an appropriate location for walls to construct transition cells. However, the results of our experiment 3 show that, when tongues first came into contact, the cells close to the junction were irregular but after further construction had taken place, we found that irregularities reduced significantly. A potential behavioural trait whereby the bees built cells suitable for an anticipated uniting of one tongue edge to another appears unnecessary as our hypothesis that size and angles are balanced across adjacent cells provides a sufficient explanation. Haphazard contact of misaligned cells created highly irregular cells, but continued construction and optimisation of each cell led to reduced cell-to-cell disparity eventually creating the conditions, as found by Smith et al, of a gradual progression of geometries across a tongue junction.

Through these experiments we have examined the response to conditions that occur during the later stages of cell construction. The results show how the interaction between two or more cells in contact with each other equilibrates so that cell walls are of equal length, corner angles settle at 120° and, emergent from these features, cells become hexagonal prisms with rhombic bases. In conclusion, *A. mellifera* strive to build cells that conform to an ideal of a cylindrical tube with a rounded base, but the proximity of adjacent cells impose constraints and so result in a self-organised mutual compromise that results in the available space and angular being shared equally between connected cells leading to regular honeycomb.

